# Domainator, a flexible software suite for domain-based annotation and neighborhood analysis, identifies proteins involved in antiviral systems

**DOI:** 10.1101/2024.04.23.590562

**Authors:** Sean R. Johnson, Peter Weigele, Alexey Fomenkov, Andrew Ge, Anna Vincze, Richard J. Roberts, Zhiyi Sun

## Abstract

The availability of large databases of biological sequences presents an opportunity for in-depth exploration of gene diversity and function. Bacterial defense systems are a rich source of diverse, but difficult to annotate genes with biotechnological applications. In this work, we present Domainator, a flexible and modular software suite for domain-based gene neighborhood and protein search, extraction, and clustering. We demonstrate the utility of Domainator through three examples related to bacterial defense systems. First, we cluster CRISPR-associated Rossman fold (CARF) containing proteins with difficult to annotate effector domains, classifying most of them as likely transcriptional regulators and a subset as likely RNAses. Second, we extract and cluster P4-like phage satellite defense hotspots and identify an abundant system related to Lamassu phage defense systems. Third, we integrate a protein language model into Domainator and use it to identify restriction enzymes with low homology to known reference sequences, validating the activity of one example in-vitro. Domainator is made available as an open-source package with detailed documentation and usage examples.

## Main Text

The rapid growth of publicly available biological sequence databases provides scientists with unprecedented opportunities to explore biological sequence diversity and co-association across virtually any range of phylogenetic and evolutionary distance. For example, multiple sequence alignments including huge numbers of homologous sequences produced by metagenome sequencing efforts around the world have been critical to the training of large-scale models for protein structure prediction^1,2^. For the individual researcher, however, more tools are needed to access the biological patterns and meanings that can be found in biological sequence space^3^. Two critical limitations hinder the ability to extract human-readable patterns necessary for the formation of testable hypotheses. First, is accurate annotation of genes at the level of functional domains, and second is the availability of tools for the automated retrieval and summarization of gene sequences within their genomic context beginning from queries utilizing combinations of functional domains of interest.

Gene annotation traditionally has been limited by approaches that use a one-to-one mapping of a known annotation onto a query sequence, for example, by transferring the annotation of the best scoring hit in a BLAST^4^ search to the query, an approach that is highly vulnerable to mis-annotation and which unfortunately can be propagated throughout databases well beyond their original scope^5^. Sequence profile-based approaches, such as HMMER^6^, together with community curated models of protein functional domains such as Pfam^7^, KOFam^8^, CAZy^9^, NCBI PGAP^10^, and others greatly mitigate this problem. Accurate domain level annotations can not only help provide consistent description of gene function, putative or otherwise, but it can also aid in the prediction of unknown functions through the co-association of functional domains within a single gene product. A clear example can be found in the polymorphic toxin systems^11^ which utilize a single polypeptide composed of a conserved carrier domain followed by variable payload effector domain for delivery via extracellular secretion machinery: new toxin functions can be identified through association with the more readily identifiable carrier domain. In another example, Lutz et al.^12^ discovered novel modification dependent DNA endonucleases by searching for PUA RNA epitranscriptomic “reader” domains fused to HNH or PD-(D/E)XK catalytic deoxyendonuclease domains. These two examples show how accurate domain annotation coupled with observation of inter-gene domain associations can be utilized to help identify gene function.

Similarly, genome context can be used to infer potential gene function by observing what other genes are in the surrounding “neighborhood” of the gene in question. Genome contextual analysis takes advantage of a general phenomenon where genes whose products function together in a biosynthetic pathway, or in a macromolecular complex, tend to cluster together at a genomic locus. Functional gene clustering is especially pronounced in bacteria and archaea and may be an adaptation for co-regulation of genes or driven by lateral gene transfer since genes that function as a “team” are more likely to provide a fitness benefit when they are transferred together into a naive genome. Regardless of the origins of gene clustering, the phenomenon can be taken advantage of when attempting to find new gene functions. If, for example, a gene encodes an enzyme in the biosynthetic pathway producing an antibiotic or other cellular metabolite, then the neighboring genes may encode enzymes either producing a substrate for that enzyme or consuming the product of the enzyme in question. The co-association of genes across multiple genomes implies that the association is under selection and therefore can be taken as further evidence of a functional interdependence. Publicly available tools enabling the retrieval of sets of genes based on their proximity include the EFI-Genome Neighborhood Tool^13^, available through the Enzyme Function Initiative, and antiSMASH^14^, a tool for identifying secondary metabolite biosynthesis gene clusters. However, these tools either do not permit querying of user input genomes or they are specialized to the mining biosynthetic gene clusters. What is needed are more general tools allowing a protein or functional domain to serve as an anchor query to user defined databases and returning hits with surrounding genes and annotations according to user specified distances.

More recently, positional/locus-based approaches have been used to find entirely new sets of gene functions without prior knowledge of any of the function of the individual genes recovered. This approach takes advantage of the phenomenon where a genetic locus itself is an “island” or “hotspot” for diverse collections of genes that can be loosely grouped together under a broader functional definition. For example, genome defense genes, that is genes involved in the protection of a cell from the incursion of non-self genes, such as from viruses or mobile genetic elements, can be found at conserved locations within genomes across a range of species. The composition of genes at these loci can vary widely across even closely related species, indicative of a high rate of gene loss/acquisition/exchange. The Immigration Control Region of the Enterobacteriaceae^15^ is a compelling example of such a dynamic locus. This region encodes many restriction endonuclease genes believed to protect the host genome against phages. Similarly, prophages themselves have been shown to encode hotspots of genome defense functions, presumably to confer some fitness advantage to their hosts during lysogeny by preventing infection from rival phages^16^. In both cases, the dynamic locus is bounded by conserved genes. Homology searches for genomic sub-sequences bounded by conserved “framework” genes can be used to recover sequences of diverse composition enriched for a common function. Flexible and automatable tools to annotate, search, extract, and organize intra-gene, gene neighborhood, and island/hotspot configurations are needed.

In addition to profile- and neighborhood-based methods, structure-based searches show high sensitivity in finding distant homologs based on shared protein fold between query and target^17,18^. However, for many users, these approaches may be limited by the computational resources needed to obtain structural models for input query sequence as well as the sequences in the target database. The availability of a variety of structural databases, either experimentally validated such as the PDB, or precomputed predictions of large datasets such as AlphaFold Protein Structure Database^19^ has alleviated the need for individual users to do structure prediction at scale. These databases, coupled with structure-based search tools such as Foldseek^18^, Rupee^20^, and Dali^17^ have resulted in a quantum leap in the ability to search for structural homologs of a protein of interest. But on the user side, the computational demands of de novo structure prediction for query sequences can still limit the number of searches possible. Methods for converting amino acid sequences directly to Foldseek-compatible encodings, without full atomic coordinate prediction, can greatly expand the search space and scale accessible to the users^21,22^.

We present here Domainator, an extended suite of command line software for sequence database search, genome annotation, and extraction and comparison of sub-genomic regions enabling rapid and flexible genomic contextual analyses for comparative genomics. We demonstrate the utility of Domainator through application to three examples. The examples presented follow a common pattern of search, annotate, report and cluster, analyze and apply, highlighting the flexibility and range of functionalities of Domainator along the way (Table 1, Fig. 1A, and 2A). In the first example, we show how Domainator can be used to cluster individual proteins and examine their domain composition, focusing on CRISPR-associated Rossman fold (CARF) domain containing proteins^23,24^. In the second example, we show how Domainator can be used to extract and analyze gene neighborhoods, focusing on P4-like phage satellite defense hotspots in *E. coli*^16^. In the third example we augment Domainator with a protein language model (pLM)^2,22^ enabling sensitive search with structural information to identify restriction modification (RM) systems where the restriction enzyme (RE) has low sequence identity to any previously characterized REs. We validate the endonuclease activity of one of the newly identified RE candidates using experimental assays.

**Table 1.**
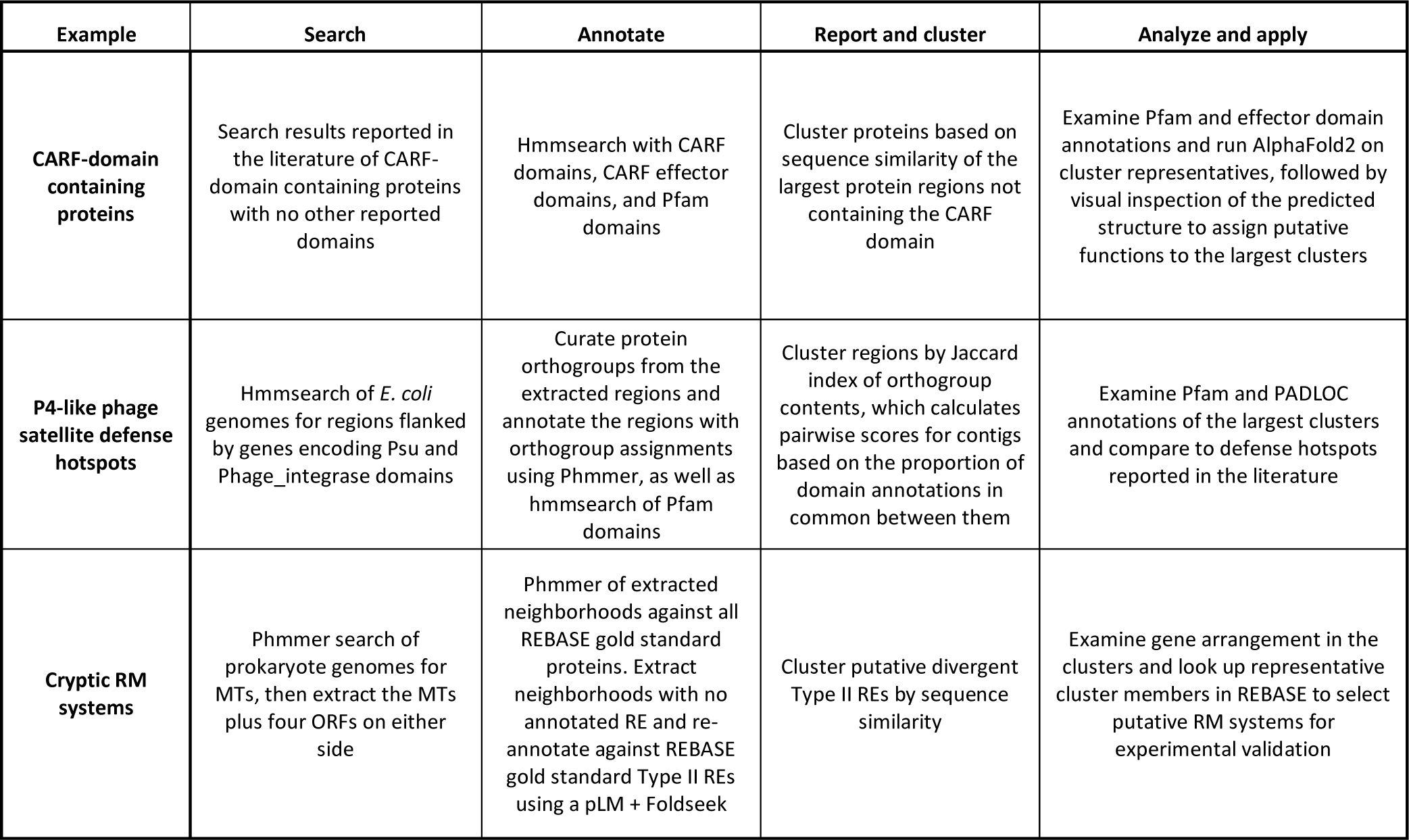
Overview of examples of practical applications of Domainator presented in this work.

| Example | Search | Annotate | Report and cluster | Analyze and apply |
| --- | --- | --- | --- | --- |
| **CARF-domain containing proteins** | Search results reported in the literature of CARF-domain containing proteins with no other reported domains | Hmmsearch with CARF domains, CARF effector domains, and Pfam domains | Cluster proteins based on sequence similarity of the largest protein regions not containing the CARF domain | Examine Pfam and effector domain annotations and run AlphaFold2 on cluster representatives, followed by visual inspection of the predicted structure to assign putative functions to the largest clusters |
| **P4-like phage satellite defense hotspots** | Hmmsearch of *E. coli* genomes for regions flanked by genes encoding *Psu* and *Phage_integrase* domains | Curate protein orthogroups from the extracted regions and annotate the regions with orthogroup assignments using Phmmer, as well as hmmsearch of Pfam domains | Cluster regions by Jaccard index of orthogroup contents, which calculates pairwise scores for contigs based on the proportion of domain annotations in common between them | Examine Pfam and PADLOC annotations of the largest clusters and compare to defense hotspots reported in the literature |
| **Cryptic RM systems** | Phmmer search of prokaryote genomes for MTs, then extract the MTs plus four ORFs on either side | Phmmer of extracted neighborhoods against all REBASE gold standard proteins. Extract neighborhoods with no annotated RE and re-annotate against REBASE gold standard Type II REs using a pLM + Foldseek | Cluster putative divergent Type II REs by sequence similarity | Examine gene arrangement in the clusters and look up representative cluster members in REBASE to select putative RM systems for experimental validation |

**Fig 1.**
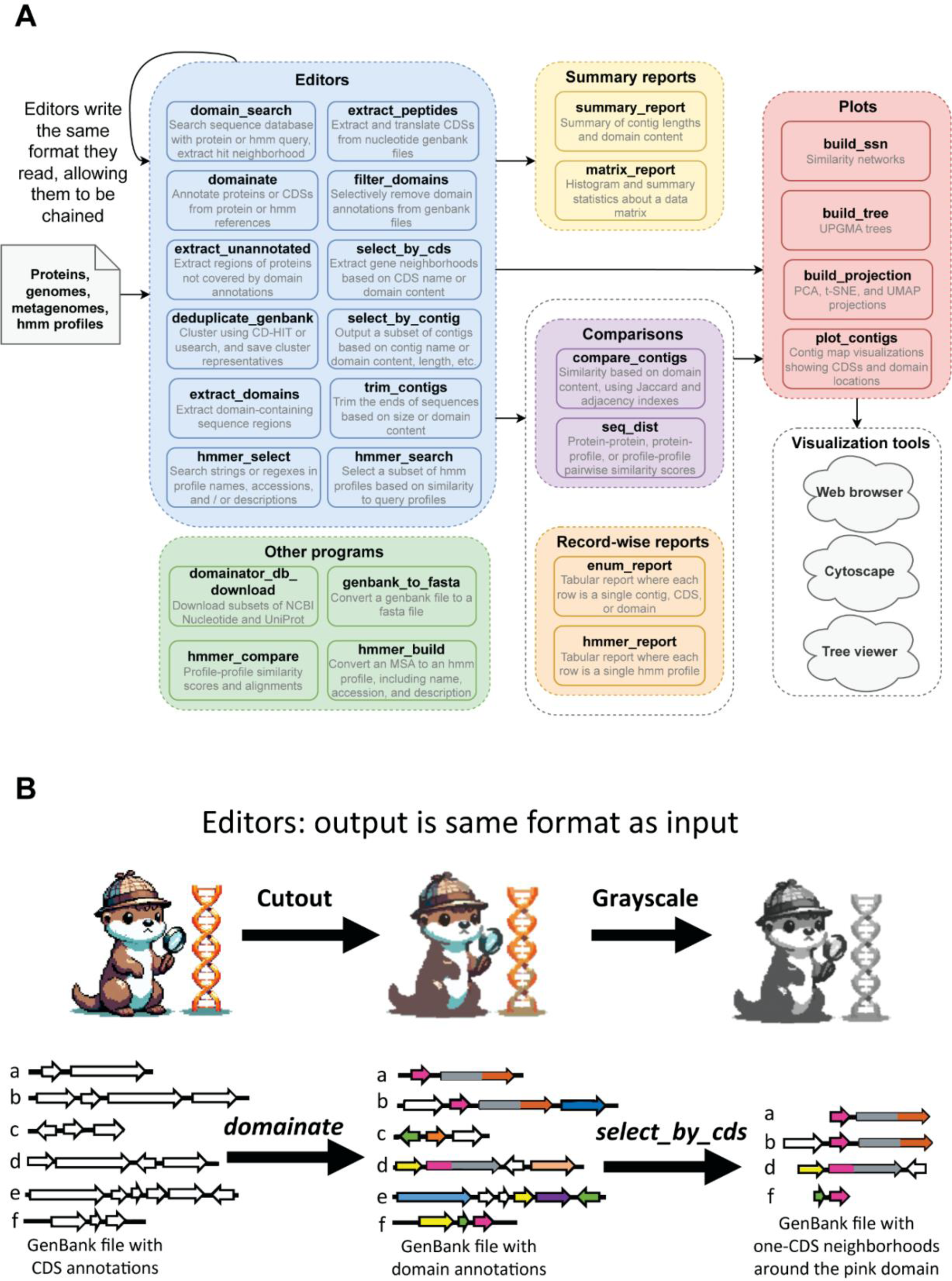
Domainator is designed around a modular architecture. (A) Examples of the different kinds of programs in Domainator (not all programs are shown). (B) Editors are a special kind of program where the input files and output files have the same format, editor programs in Domainator can apply sequential transformations to GenBank files, similar to how image editors can apply sequential transformations to image files.

## Results

### The Domainator toolkit

Domainator provides more than two dozen discrete, flexible programs that can be composed into a broad range of genome mining and comparative genomics workflows via command line or python scripting (Fig. 1A). The high degree of modularity built into Domainator stands in contrast to other genome mining tools, such as EFI ^25^, antiSMASH^14^, BiG-SCAPE/CORASON^26^, and MacSyFinder^27^, which supply dedicated end-to-end workflows for specific tasks. Another unique feature is that Domainator supports a rich set of operations on HMM-profiles, for example, subsetting of .hmm files and comparison of HMM-profiles, including the construction of profile vs. profile similarity networks and trees (Fig. 2B). The “Domain” part of the name “Domainator” derives from the key role that local (*i.e.*, subsequence) alignments play in Domainator workflows. Domainator uses the GenBank file format^28^ (https://www.ncbi.nlm.nih.gov/genbank/samplerecord/) as a carrier of both sequence and annotation data. Independence from a fixed set of sequence sources and the co-location of sequences and all their annotation data in a single file increases data portability and decreases complexity for end-users. Domainator can add functional annotations to sequences by local alignments against databases of HMM-profiles, protein sequences, or both at the same time. For example, in a single call to the *domainate* program, a set of genome or metagenome contigs can be annotated with hits to Pfam HMM-profiles^7,29^ and hits to REBASE Gold Standard protein sequences^30^ at the same time. De novo annotations derived by Domainator can be added atop pre-existing metadata in Genbank format files, but the software also provides options to filter or even replace earlier annotations.

**Fig 2.**
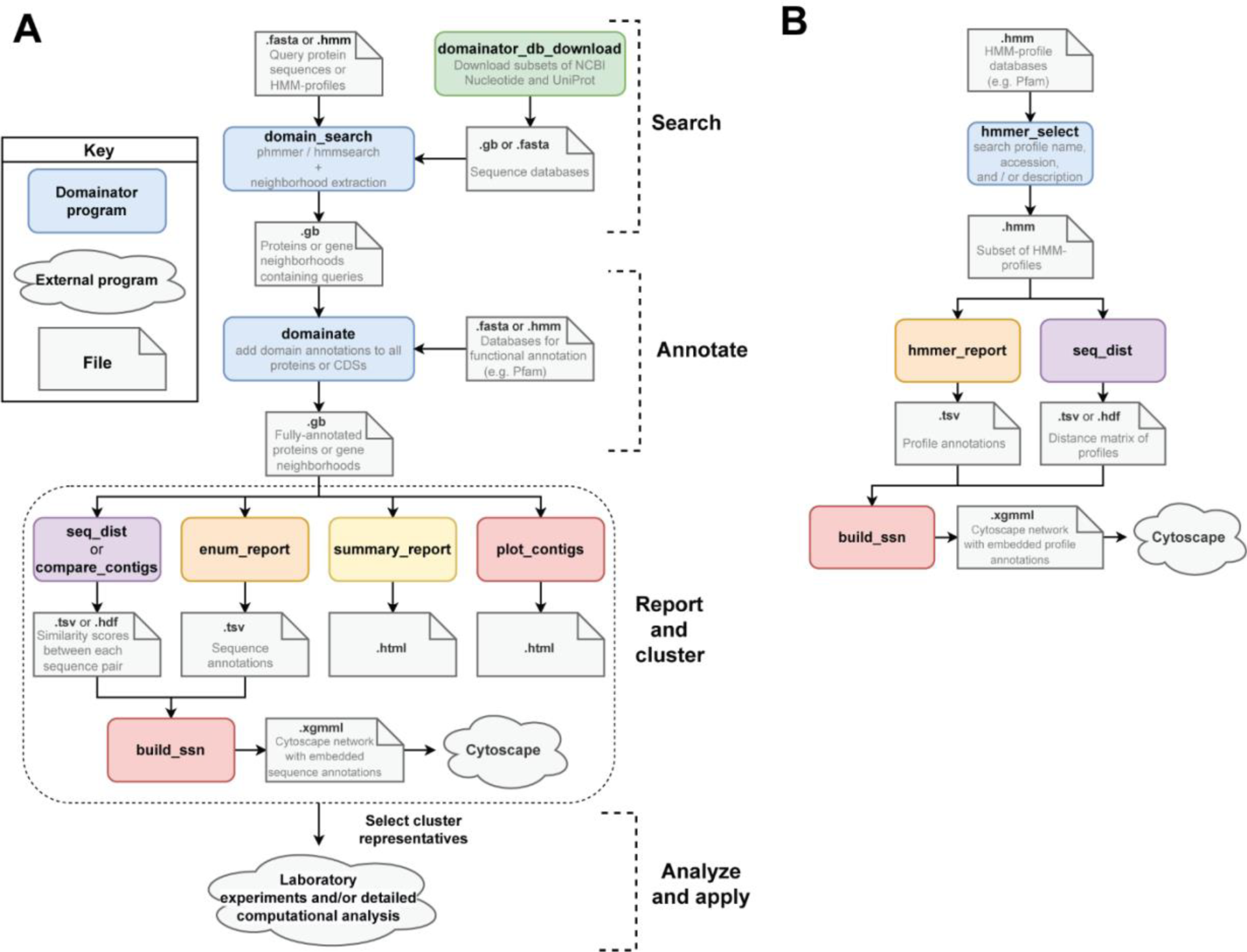
General strategy for using Domainator to annotate and compare (A) proteins and gene neighborhoods, and (B) HMM profiles.

The individual programs that make up Domainator can be roughly grouped into six categories corresponding to their general roles in genome mining and comparative genomics workflows as diagrammed in Fig. 1A. The first steps in most workflows often involve passing sequence data through one or more editors.

> **Editors** are programs whose output format is the same as their input format. Each individual editor performs a simple task, such as adding putative functional annotations to a GenBank file or extracting a subset of contigs, but they can also be combined in arbitrarily long chains to accomplish complex transformations (Fig. 1B). Examples are *domain_search*, which outputs a subset of the input sequences, based on the presence of a hit to a reference sequence or profile; *domainate*, which outputs all the input sequence but adds domain annotations based on hits to user-specified reference sequences; *deduplicate_genbank*, which performs similarity clustering using CD-HIT^31^ or USEARCH^32^ and outputs only the cluster representatives of the input sequences; and *select_by_cds*, which extracts genome neighborhoods (including sequence, features, and annotations) surrounding domains of interest.

> **Summary report** programs summarize data into graphs and statistics, for example, the number of sequences in a file, the count of each kind of domain, and the distribution of taxonomic origins of the sequences. Reports are provided in human readable format either as text displayed in the console or as HTML files.

> **Record-wise report** programs produce tab-separated files, for example where each row corresponds to a genome contig, a protein, or a domain, and columns are data such as length, taxonomy ID, domain content, etc. Record-wise reports are useful for exporting data to programs, such as Excel, which can’t read GenBank or hmm files, and they also find use as intermediary files between some programs in Domainator.

> **Comparison** programs generate pairwise score or distance matrices between proteins, contigs, or HMM-profiles. *Compare_contigs* uses the Jaccard^33^ and adjacency^26^ indexes to compare proteins or gene neighborhoods based on their domain content, whereas *seq_dist* uses local alignment scores, for example via phmmer^6^, Diamond^34^, hmmsearch^6^, or the Viterbi profile-comparison algorithm ^6,35,36^. Comparison programs output pairwise distance or score matrices that can be presented in tables or used to plot trees or similarity network diagrams.

> **Plotting** programs convert data into formats appropriate for graphical visualization, for example converting distance or score matrices and tabular metadata into trees or similarity networks which can be viewed in Cytoscape^37^ or other external visualization tools, depending on the data type. Plotting programs that take matrices as input can be also plot data from programs outside the Domainator suite. For example, *build_ssn* could be used to generate a structure similarity network by supplying it with a text file containing a table of pairwise structural similarity scores between a set of .pdb structure files, as calculated by TM-align^38^ or some other program.

Finally, there are a few **other** programs that defy categorization. These programs perform functions such as downloading data from NCBI or UniProt, converting files between formats, or generating profile-profile alignments.

The modularity of Domainator makes it straightforward to integrate the next generation of protein annotation methods into Domainator workflows, including methods pLM embeddings^2,39^. We recently reported a method^22^ to greatly improve the sensitivity of remote homology detection between proteins using the ESM-2 3B pLM^2^ fine-tuned to convert amino acid sequences directly into the Foldseek 3Di structure alphabet^18^. The fine-tuned model, ESM-2 3B 3Di, generates 3Di sequences roughly 1,000 times faster than AlphaFold2. The 3Di sequences predicted by the model perform well when used as queries and targets in Foldseek searches, outperforming both phmmer and hmmscan for sequences less than 20% identical to each other^22,40^. We integrated ESM-2 3B 3Di into the *domainate* program, allowing annotation of protein sequences by on-the-fly conversion to 3Di sequences followed by searches against Foldseek databases.

### Comparing CRISPR-associated Rossman fold containing proteins

CARF containing proteins are ancillary proteins, commonly associated with type III CRISPR-Cas systems^24,41–43^ (Fig. 3A). In some type III systems, the Cas10 subunit has a cyclic oligoadenylate (cOA) synthase activity that becomes active when the type III complex binds to target RNA. CARF containing proteins in these systems typically have two domains, the CARF domain and an effector domain. Binding of the cOA signaling molecule by the CARF domain induces homo-dimerization and activation of the effector domain. The effector domain is highly variable, but often functions as a non-specific nuclease. Activation of the effector domain provides an additional layer of immunity to the targeted nuclease activity of the CRISPR-Cas system, in some cases slowing down host growth or killing the host in an abortive infection process. Characterization of the variable domains fused to the conserved CARF domains can lead to discovery of new defense-related enzymatic activities.

**Fig 3.**
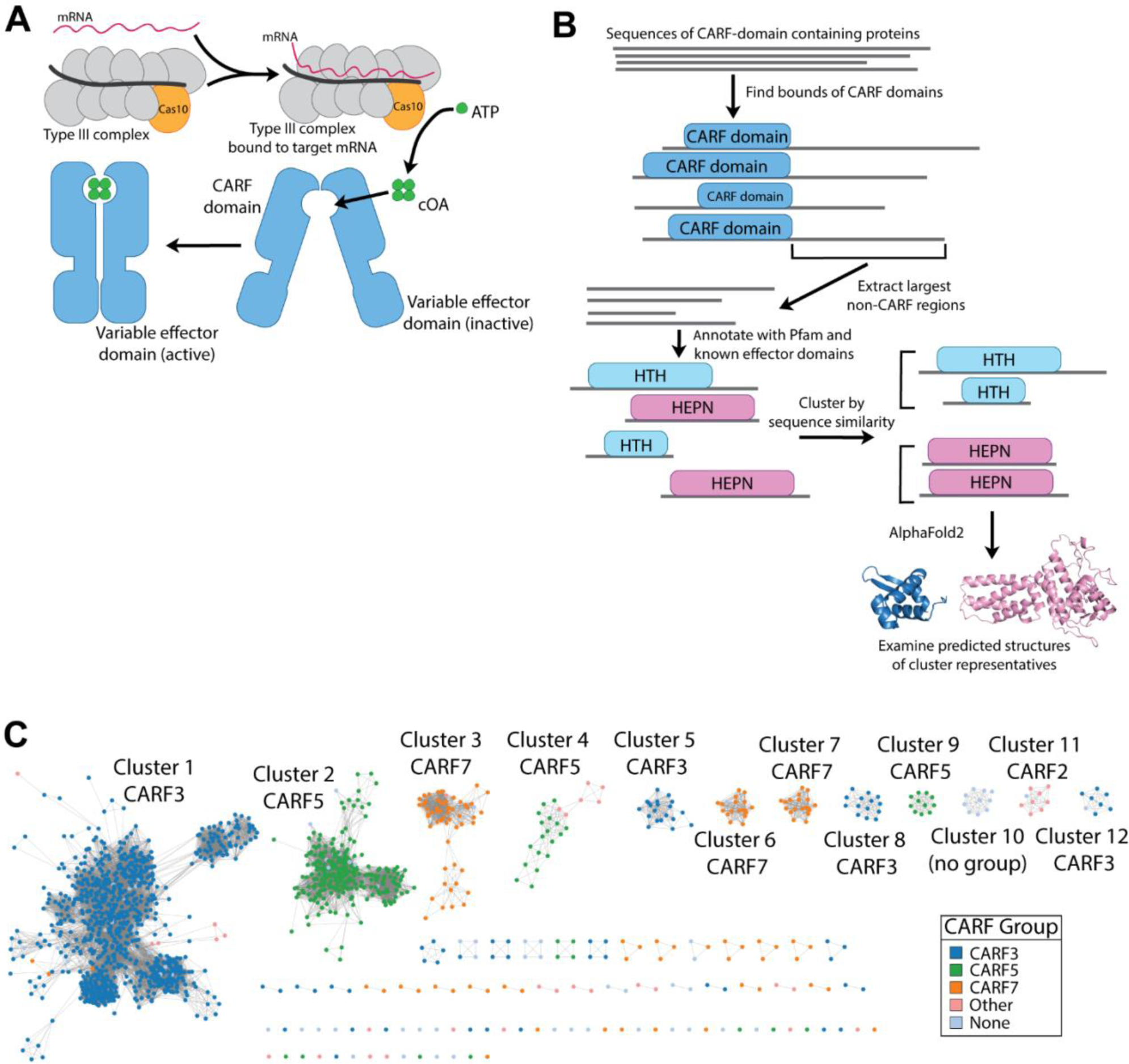
Annotating variable effector domains of CARF-domain containing proteins. (A) Schematic of effector activation. (B) Overview of bioinformatics strategy. (C) Sequence similarity network of effector domains, colored based on their associated CARF domain.

To extend the analysis of CARF domain containing proteins recently catalogued by Makarova, Koonin, and colleagues^24^, we used Domainator in a workflow assigning putative annotations to fused domains left unannotated in the earlier work (Figs. 3B,C and S1, Table 2). Searching through a database of 13,116 completely assembled archaeal and bacterial genomes, these authors compiled a list of 6,665 CARF-domain containing proteins, sorting them into 25 clades, and annotated the fused domains. In 1,844 of the CARF-domain containing proteins, they identified a single CARF domain and no additional domain annotations.

**Table 2.** CARF effector cluster annotations.

| Cluster | Count | CARF Clade (count) | Putative annotation | Evidence |
| --- | --- | --- | --- | --- |
| 1 | 916 | CARF3 (886), CARF_m5 (5), CARF7 (4), CARF1 (1) | Transcriptional regulator | Pfam annotations, apparent wHTH domain in AlphaFold2 structure. |
| 2 | 282 | CARF5 (273) | Transcriptional regulator | Apparent wHTH domain in AlphaFold2 structure. |
| 3 | 59 | CARF7 (58) | Transcriptional regulator | Apparent wHTH domain in AlphaFold2 structure. |
| 4 | 24 | CARF5 (18), CARF1 (4), CARF_m13 (2) | Transcriptional regulator | Apparent wHTH domain in AlphaFold2 structure. |
| 5 | 20 | CARF3 (20) | Transcriptional regulator | Apparent wHTH domain in AlphaFold2 structure. |
| 6 | 19 | CARF7 (18) | CARF7_DUF2103 | Weak hits to CARF7_DUF2103 profile |
| 7 | 19 | CARF7 (19) | Transcriptional regulator | Apparent HTH domain in AlphaFold2 structure. |
| 8 | 14 | CARF3 (14) | Transcriptional regulator | Apparent wHTH domain in AlphaFold2 structure. |
| 9 | 11 | CARF5 (11) | CARF7_DUF2103 | Contains apparent HTH domain, C-terminal domain is a weak hit to CARF7_DUF2103 profile |
| 10 | 10 | None | Unknown |  |
| 11 | 9 | CARF2 (8) | HEPN nuclease | Contains conserved HEPN RNase active site residues RNXXXH |
| 12 | 8 | CARF3 (7) | Transcriptional regulator | Pfam annotations, apparent wHTH domain in AlphaFold2 structure. |

For each of these CARF-only proteins, we identified the footprint of the putative CARF domains and extracted the largest region not covered by those domains, provided it was 50 amino acids or longer, producing a list of 1,524 protein fragments. We annotated the fragments using hmmscan against Pfam (v 36)^7^ and profiles built from CARF effector domain alignments reported in the earlier work^24^. We built a sequence similarity network from the fragments (Fig. 3C), grouping them into 94 homology clusters. Within each cluster, almost all the members are fusions to CARF domains of the same clade. This co-occurrence is notable because we clustered based on just the fusion domain rather than the CARF domain or entire protein, suggesting coevolution. The top twelve homology clusters contained 1,391 of the 1,524 protein fragments, with 916 fragments belonging to the largest cluster (Table 2, Table S1). We assigned a putative functional annotation to each cluster (Table 2) using Pfam and effector domain annotations, CARF clade, and examination of AlphaFold2 predicted structures^1,44^ (Fig. S2, Table S2).

AlphaFold2 structures of randomly selected examples of 9 out the 12 clusters contain apparent Helix-Turn-Helix (HTH) or winged Helix-Turn-Helix (wHTH) motifs, indicating that they likely bind DNA and may serve as transcriptional regulators. In some cases, the DNA binding motif comprises the entire non-CARF region. In the case of clusters 2, 3, and 9, additional protein sequence may confer catalytic activity or some other function. Clusters 6 and 9 appear to be divergent members of the previously catalogued CARF7_DUF2103 domain family^24^. Cluster 11 appears to contain HEPN nucleases^45^, non-specific RNAses that are one of the most common effector domains found fused to CARF domains. We were unable to determine a putative annotation for cluster 10.

The example of CARF effector domain analysis demonstrates some of the capabilities of Domainator to extract, cluster, and annotate individual proteins and protein fragments. In our next example, we show how Domainator enables the discovery of multigene defense hotspots through the same paradigm of search, annotate, cluster, and analyze.

### Comparing P4-like phage satellite defense hotspots in *E. coli*

Diverse defense systems in bacteria are often found in bacterial genomes as variable defense islands, hotspots, or cassettes, flanked by conserved “framework” genes. We use the example of P4-like satellites (Fig. 4A), previously catalogued by Rousset et. al (2022)^16^, to demonstrate the capability of the Domainator suite for identifying, extracting, and analyzing variable defense regions. Using a series Domainator commands (Fig. S3), we started from a set of 2,718 *E. coli* genomes and extracted 910 regions flanked by Psu and Phage_integrase Pfam (version 36) domains^7^, between 1 and 9 CDSs in size. We then produced a tabular summary of each instance of the hotspot (Table S3), and a graphical neighborhood similarity network (Fig. 4B) based on the pairwise Jaccard indexes^26^ of orthogroup contents (Table S4), grouping the hotspots into clusters with shared orthogroup contents. The Jaccard index is calculated by dividing the number of unique orthogroups in common between two neighborhoods by the total number of unique orthogroups in the two contigs. A limitation of similarity networks is that they do not show the relationship between clusters. To get a more granular view of cluster relationships, Domainator can also produce Jaccard index trees, by Jaccard distance as 1 - Jaccard index, followed by calculating an unweighted pair group method with arithmetic mean (UPGMA) tree (Fig. S4), from the pairwise Jaccard distances.

**Fig 4.**
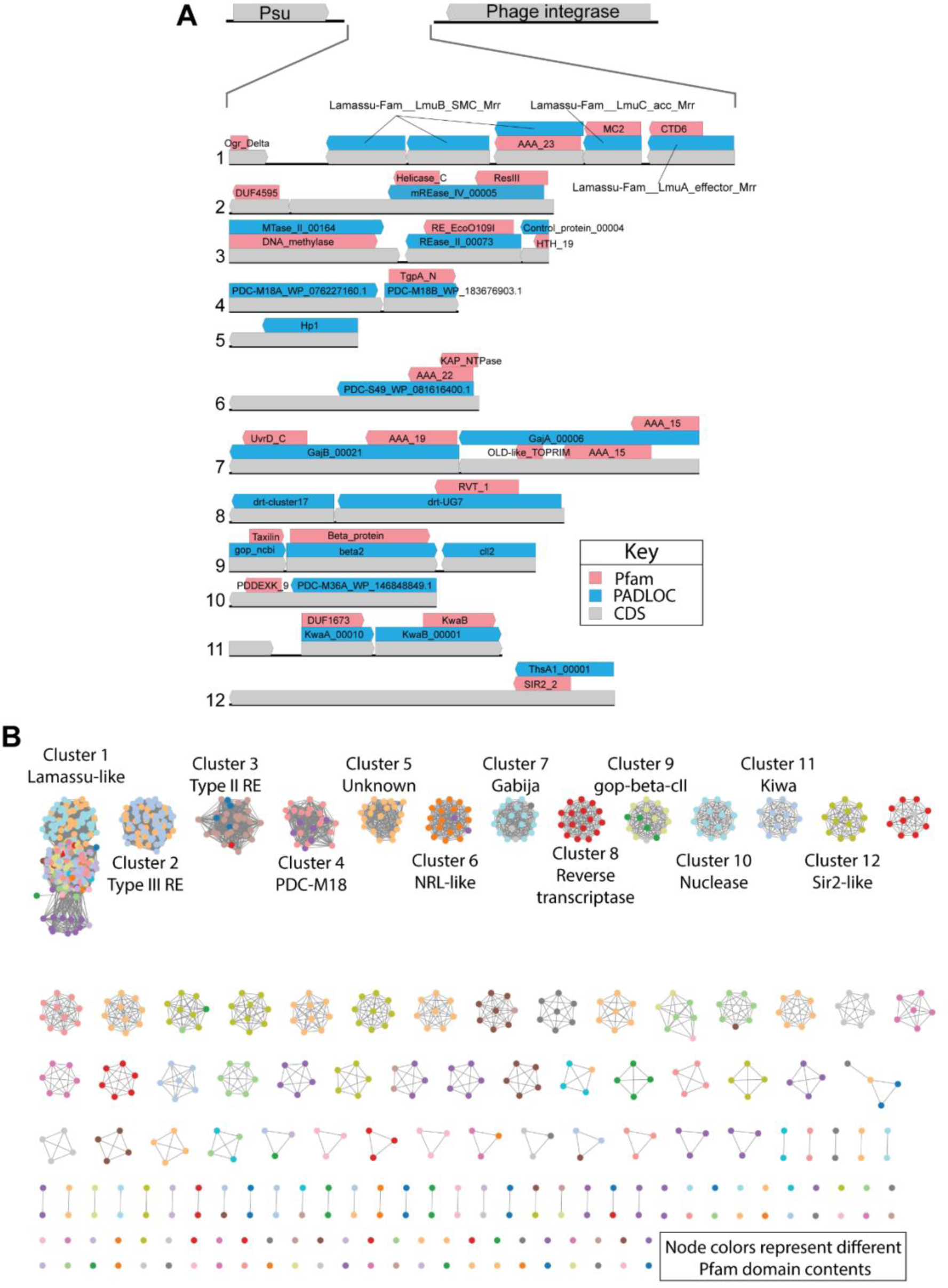
Identifying and categorizing P4-like phage satellite defense hotspots in *E. coli*. (A) P4-like phage satellite defense hotspots are highly variable regions of 1 to 9 genes found between Psu and Phage integrase genes. Pfam and PADLOC annotations of representatives from the top 12 clusters are shown. (B) A similarity network of P4-like phage satellite defense hotspots, edges are drawn between clusters with orthogroup content Jaccard index of greater than 0.7, colors represent different Pfam domain arrangements within the hotspot.

The results of our search were broadly consistent with the previously reported results^16^ (Table 3), despite our use of a different set of starting genomes and different analysis methods. Of the top 12 most frequent defense systems found through our analyses, 9 were among the top 12 most abundant systems reported by Rousset et al., and 11 here were among their top 20. These systems include, restriction endonucleases^46^, the Kiwa^47^, and Gabija^47,48^ systems, among others. Surprisingly, the most frequent system in our analysis was not previously described, but has some homology to Lamassu systems^49,50^, based on annotation with the PADLOC database^51^. Upon further investigation, these hotspots were not from P4-like phages, but apparently from satellites derived from other Caudoviricetes phages, as determined by Blast search of the satellites and their flanking regions against viral genomes in NCBI nr. Furthermore, their flanking Psu sequences had much lower identities to the reference Psu sequence (GenBank: WP_000446153) than did Psu sequences flanking other common clusters. Psu sequences from the most abundant cluster had an average of 31% amino-acid sequence identity to the reference P4 Psu, whereas Psu sequences from other clusters averaged 98% or 99% identity to the reference (Tables 3 and S5).

**Table 3.**
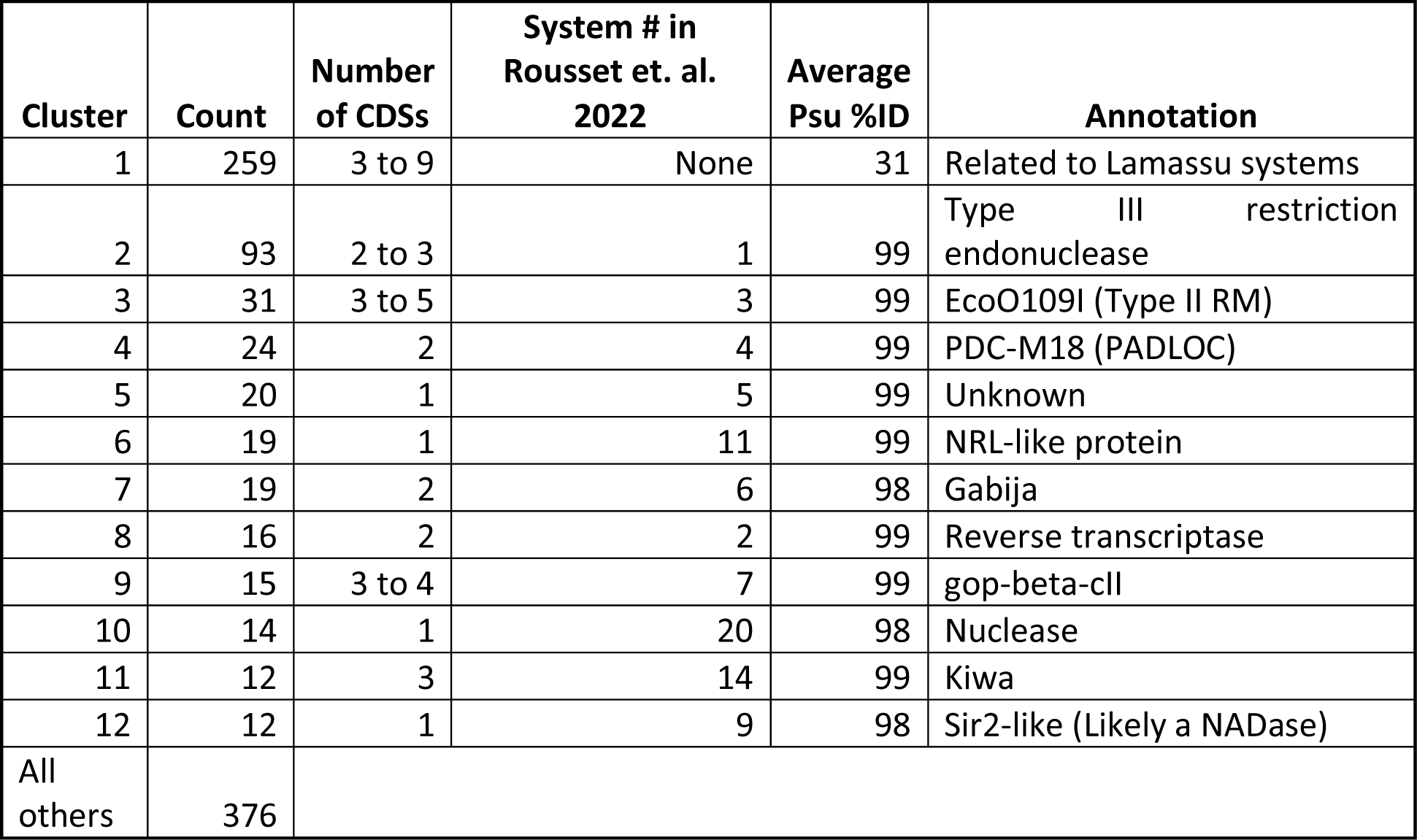
P4-like phage satellite defense hotspot annotations. Psu % identity is relative to the P4 phage reference sequence (GenBank: WP_000446153).

The example of extracting and comparing P4-like phage satellite defense hotspots demonstrates the power of Domainator in the analysis of gene neighborhoods, enabling the discovery of variable defense regions, and uncovering potential new defense systems from Caudoviricetes, distantly homologous to the previously described defense hotspots in P4-like satellites. In our next example, we augment Domainator with a protein language model to assign useful functional annotations to proteins even deeper into the twilight zone of similarity to well-characterized reference sequences.

Identification of cryptic restriction modification systems using a protein language model Restriction endonucleases (REs) are enzymes that cut foreign DNA upon entry into a bacterial cell and thus play an important role in bacterial defense against bacteriophages^52^. As a whole, restriction endonucleases are highly diverse, but can be categorized according to Types I-IV. Type I, II, and III enzymes utilize DNA methylation patterns established by methyltransferases (MTase) in the host cell to distinguish self from non-selfDNA. These two enzymatic activities are dependent on each other for a functioning restriction-modification (RM) system. Methylation and cleavage are specific to a shared DNA sequence motif, and methylation of DNA blocks cleavage of DNA by the cognate RE. In this way, the MT protects the bacterial genome from cleavage by its own RE. Foreign DNA, such as bacteriophage DNA originating from an environment lacking an appropriate MT, will be susceptible to being cut by an RE. The methyltransferase (MT) and corresponding restriction enzyme (RE), are typically encoded by genes within four open reading frames (ORFs) of each other ^30,53^ (Fig. 5A). MTs tend to exhibit slower evolution and sequence divergence than REs^30^, perhaps because of their more complex chemistry and the weaker selection pressure in the absence of phage threats. The implication is that MTs are more easily detectable than REs by primary sequence search such as BLASTP^54^ or phmmer^6^. Therefore, it can be expected that bioinformatic discovery of RM systems using sequence homology search methods may fail to identify RM systems where the MT is detectable by sequence search, but a neighboring RE gene has diverged too far from the reference sequences to be detectable.

**Fig 5.**
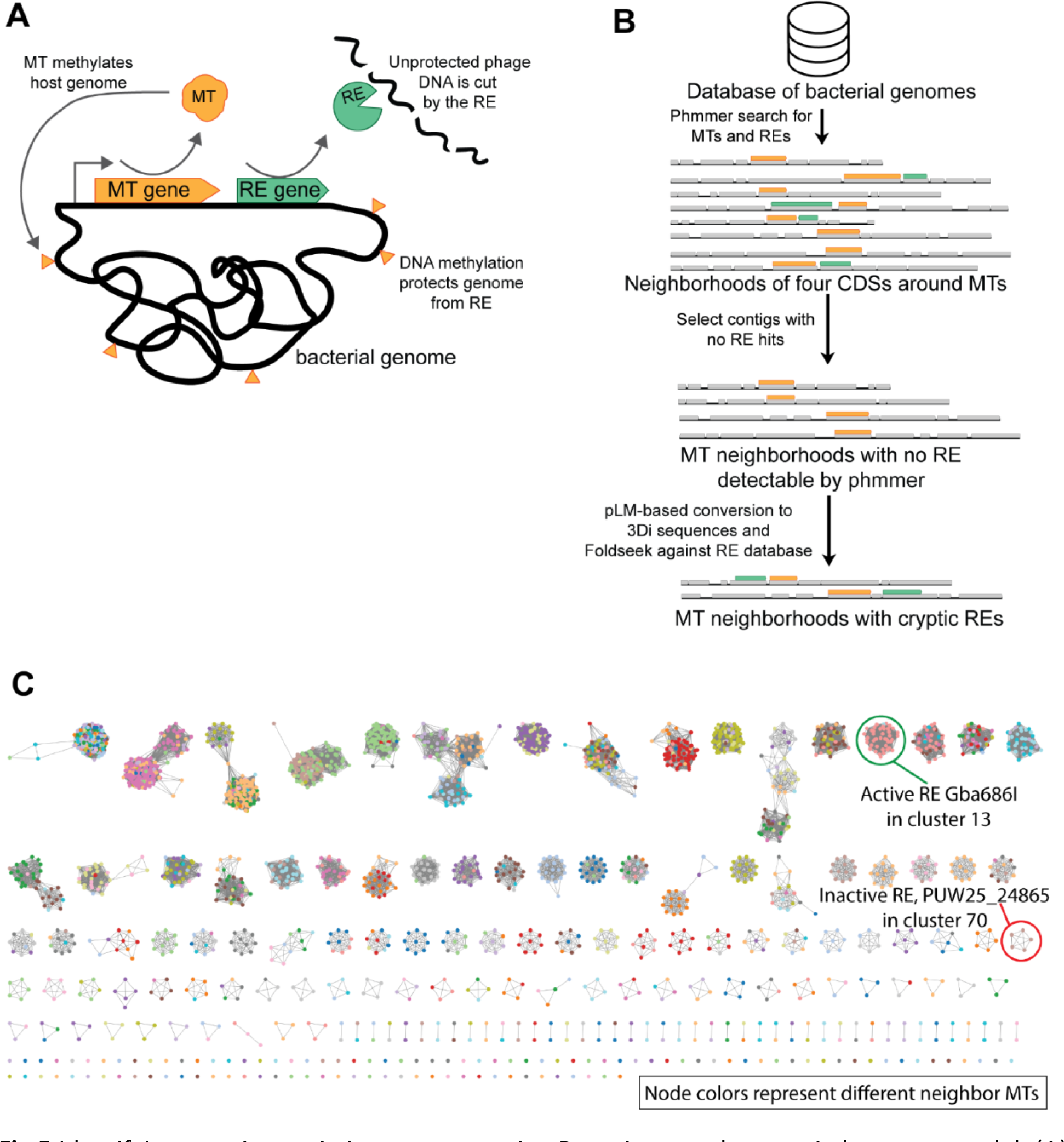
Identifying cryptic restriction enzymes using Domaintor and a protein language model. (A) Schematic of how Type II RM systems protect bacteria from phages. (B) Overview of bioinformatics strategy. (C) Sequence similarity network based on similarity of predicted REs, and colored based on the best reference MT hit to the neighbor MT.

We leveraged the REBASE Gold Standards set of reference MTs, REs, and associated proteins^30^ together with the Domainator interface to phmmer, Foldseek^18^, and neighborhood extraction functionalities to identify RM systems with divergent REs (Figs. 5B and S5). We started by downloading a set of high-quality genomes from GenBank^28^ and extracting 205,039 neighborhoods containing MT phmmer hits plus four CDSs on either side of the query match. Filtering out those neighborhoods containing REs detectable by phmmer, resulted in 124,853 putative orphan MTs and their surrounding features. We then used the ESM-2 3B 3Di^22^ pLM to recode the sequences into a tertiary structure alphabet and used Foldseek^18^ to find structural matches to Type II REs within these neighborhoods, yielding 2,628 contigs containing putative cryptic RM systems (Table S6). We then generated a sequence similarity network of the REs encoded in these systems (Fig. 5C), and manually inspected representative systems selected from each cluster. We selected two candidate RM systems for experimental validation, prioritizing candidates where the putative RE is immediately adjacent to the putative MT and where the RE is absent from the REBASE^30^ entry for the corresponding genome.

The candidate restriction endonucleases selected for activity screening were: locus_tag H1R19_03490 (GenBank protein QMT02248.1) from *Gordoniaceae jinghuaiqii* str. zg-686 (GenBank nucleotide: CP059491), and locus tag PUW25_24865 (GenBank protein: WDI02380.1) from *Paenibacillus urinalis* (GenBank nucleotide: CP118108). A schematic outlining the construction of templates for *in vitro* protein expression, beginning with synthetic gene blocks, is shown Fig. 6A. Following expression using the PurExpress *in vitro* transcription/translation system, one-, three-, and nine-fold dilutions of reactions were incubated for 1 hour in 1X CutSmart buffer with 1 µg of DNA substrates: 42 kb long phage λ genomic DNA free of Dam methylation and a 22 kb plasmid pAd2BsaBI, purified from a *dam*^+^ *E. coli* strain. The *E. coli dam* gene encodes a DNA methyltransferase (Dam) modifying N6-adenine in the context of a GATC sequence. DNA from the cleavage assay reactions was resolved and visualized by agarose gel electrophoresis and ethidium bromide staining. As seen in Figure 6B, reactions expressing H1R19_03490 from *Gordonia* showed specific cleavage activity towards unmethylated λ DNA (Fig. 6B). As a result of the confirmed activity, H1R19_03490 has been renamed Gba686I, according to accepted nomenclature^55^. Neither pAd2BsaBI nor methylated (*dam*^+^) λ DNA was digested by this enzyme. Reactions expressing PUW25_24865 from *P. urinalis* did not contain detectable activity in our assay (Fig. 6B). The digestion pattern produced by Gba686I reactions was identical to that produced by BamHI (Genbank: QDP93514.1), indicating that Gba686I recognized the sequence GGATCC. However, unlike BamHI, the Gba686I is blocked by Dam methylation (Fig. 6C). We were unable to detect any amino-acid sequence similarity between Gba686I and BamHI.

**Fig 6.**
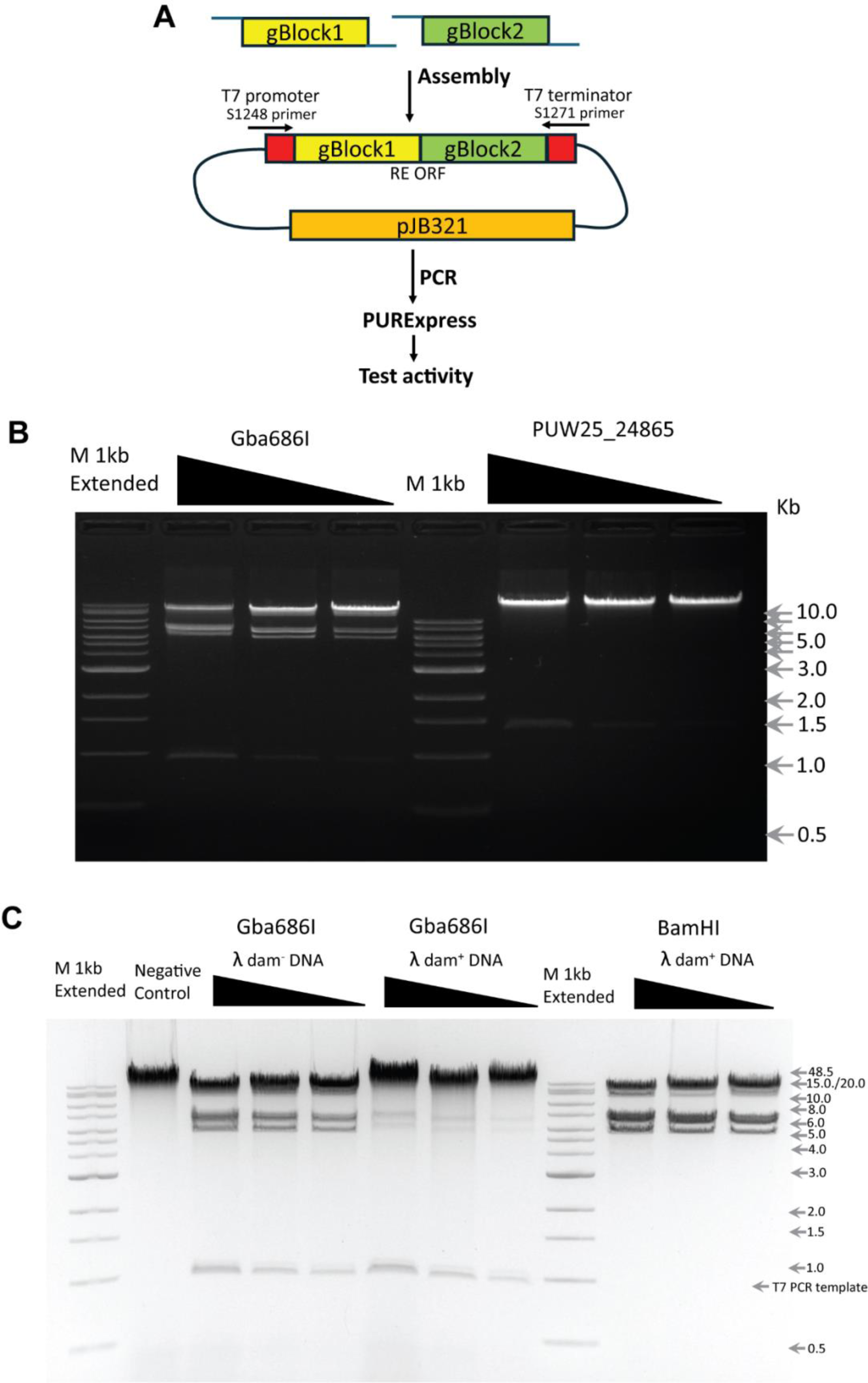
Putative RE candidate *in vitro* assay. (A) The workflow of expression experiments, including gene assembly, preparation of expression templates by PCR, and *in vitro* expression in PURExpess system. (B) Endonuclease activity assay on λ dam-DNA substrate, where Gba686I shows specific cleaveage, but PUW25_24865 is inactive. (C) Gba686I produces the same cleavage pattern as BamHI, which recognizes a GGATCC motif but, in contrast to BamHI, this isoschizomer is sensitive to Dam methylation at the GATC sequence nested within RE cleavage site. Note the presence of a DNA band at 1 kb corresponding to the expression template and not derived from the cleavage of the input substrate DNA.

## Discussion

In this work, we introduce the Domainator software suite and show how it can be used to find, extract, and cluster proteins and gene neighborhoods. We highlighted three examples of how Domainator can be used and extended to answer different kinds of questions in the context of bacterial defense systems. The three examples all follow the general pattern of search, annotate, report and cluster, analyze and apply (Fig. 2A), but instantiate that pattern using different combinations of Domainator programs to study various aspects of diverse biological systems (Table 1).

In the first example, we clustered CRISPR-associated Rossman fold containing proteins and added putative annotations to examples that were previously difficult to annotate, identifying a subset that appear to contain HEPN RNAse domains. In the second example, we extracted and clustered P4-like phage satellite defense hotspots from *E. coli*, identifying a distantly related defense hotspot with apparent homology to Lamassu systems. In the final example, we integrated a neural network pLM into Domainator and used the new functionality as part of a workflow to identify divergent restriction enzymes (RMs) that were undetectable using primary sequence-based searches. We verified the sequence-specific endonuclease activity of one of these RMs in-vitro.

The power and flexibility of Domainator was enabled by several key design choices. One key was to use the GenBank file format as the primary carrier of sequence information and annotations. The main advantage of using GenBank files as primary carriers of information is that it makes data at each intermediate step of the analysis self-contained and portable. The biggest tradeoffs of using GenBank files is that search speed can suffer due to a lack of pre-indexing of the GenBank files and the slowness of parsing GenBank files in python.

Another key choice was the separation of editor, comparison, reporting, and plotting tools into distinct scripts with modular, interchangeable and, where possible, portable input and output formats. While this separation of functions increases the complexity of using Domainator compared to other software packages for gene neighborhood and protein annotation, it gives the user much greater flexibility in composition of analysis workflows. One way we take advantage of this flexibility is by using the *build_ssn* program to build not only sequence similarity networks, but also neighborhood domain composition similarity networks, and to incorporate into those networks’ annotations produced by *enum_report* and by external sources of tab-separated data. Another advantage of the modularity is that it makes it straightforward to integrate additional functionality into Domainator, for example the pLM.

Getting started with Domainator is easy. Installation can be done in a conda environment using conda install. The available programs are detailed in a README file and additional help and usage examples can be called by executing the program with the `-h` (help) option. All a user needs are the nucleotide or protein sequences they want to filter or annotate in fasta or GenBank format, and protein sequences or HMM profiles to use as references for the annotation in fasta or hmm format. The included script, *domainator_db_download*, facilitates downloading sequences from NCBI or UniProt, with options for filtering by taxonomic origin of the sequences. Curated HMM profiles can easily be obtained from InterPro^7,29^ (https://www.ebi.ac.uk/interpro/), NCBI PGAP^10^ (https://ncbiinsights.ncbi.nlm.nih.gov/tag/hidden-markov-models-hmm/), KEGG KOfam^8^ (https://www.genome.jp/tools/kofamkoala/), or various more specialized sources, such as CAZY/dbCAN^9,56^ (https://bcb.unl.edu/dbCAN2/) for carbohydrate-active enzyme, or PADLOC^51^ (https://github.com/padlocbio/padloc-db) or DefenseFinder^57^ (https://github.com/mdmparis/defense-finder-models/) for proteins related to defense and conflict systems. Documentation, tutorials, and example workflows are available on the Domainator GitHub page (https://github.com/nebiolabs/domainator).

The three examples presented in this work just scratch the surface of what is possible using Domainator. Other potential applications include, but are not limited to, identification, extraction, and analysis of novel domain fusions and multi-domain proteins, prophages, natural product biosynthetic gene clusters, serotype clusters, pathogenicity islands, phage inducible chromosomal islands, and other mobile genetic elements, such as integrative conjugative elements (ICEs). Furthermore, Domainator seamlessly annotates and extracts intron-containing genes, so it can be used to study eukaryotic genomes as well as eubacterial and archaeal genomes. We intend to continue adding new functionality in the coming years as we use it in our own research projects. By making Domainator available free and open source, we hope that it will serve as a useful tool for the life sciences community, and we invite suggestions and contributions for its continued improvement.

## Materials and Methods

### Domainator Implementation

Domainator is written in Python and built on top of some critical dependencies, including, but not limited to BioPython^58^, HMMER3^6^ via PyHMMER^59^, Numpy^60^, SciPy^61^, Pandas^62^, CD-HIT^31^, DIAMOND^34^, Prodigal^63^ via Pyrodigal^64^, Foldseek^18^, ESM2^2^. Sequence similarity network generation draws inspiration from and uses the sequence similarity score proposed by EFI^65^. *Compare_contigs* uses the Jaccard^33^ and adjacency^26^ indexes to compare proteins or gene neighborhoods based on their domain content, whereas *seq_dist* uses local alignment scores, for example via phmmer^6^, Diamond^34^, hmmsearch^6^, or the Viterbi profile-comparison algorithm^6,35,36^.

### Key formulas

#### EFI alignment score for calculating sequence similarity^65^

EFI score = Log_10_(2^-1 · bit score of local alignment^ · (seq_1 total length · seq_2 total length))

Where the bit score can come from any local alignment algorithm, in our case, Diamond^34^. The EFI score is roughly proportional to the negative logarithm of the E-value.

#### Jaccard index for calculating similarity of domain content^33^

Jaccard index = number of unique domain annotations shared between the two contigs / total number of unique domain annotations present in either contig

### Examples

Computational workflows for the three examples are outlined in the results section above, and including as detailed flow charts in Figs. S1, S3, and S5. Code for replicating all three examples is available on GitHub (https://github.com/nebiolabs/domainator_examples). Sequence similarity networks were visualized with Cytoscape^37^, structure predictions were performed with AlphaFold2^1^ via local Colabfold^44^ and rendered in Pymol^66^.

### DNA assembly, PCR amplification and PurExpress analysis of putative restriction endonucleases

Two candidate ORFs with codon optimization for *E. coli* expression sequences for putative restriction endonuclease (RE) genes, locus_tag H1R19_03490 (GenBank protein QMT02248.1) from *Gordoniaceae jinghuaiqii* str. zg-686 (GenBank nucleotide: CP059491), and locus tag PUW25_24865 (GenBank protein: WDI02380.1) from *Paenibacillus urinalis* (GenBank nucleotide: CP118108) were assembled into the pJB321 (Julie Beaulieu, unpublished NEB, MA) vector under control of T7 promoter of with pJB321_Nde_68R and pJB321_BamHI_68F primers (Table S7) using 1 or 2 synthetic gBlocks (IDT, IA). The sizes of the gBlocks were validated on 1% DNA agarose gel (Fig. S3). The synthetic gBlocks were assembled into pJB321 by NEBuilder HiFi DNA assembly Master mix (E2621, NEB, MA) and the resulting T7 PCR DNA templates were amplified with Q5 “Hot Start” DNA polymerase (M0543, NEB, MA) and T7 forward S1248 and T7 reverse S1271 primers (Table S7) for *in vitro* protein synthesis in PurExpress system (E6800, NEB, MA). PCR products were purified with Monarch PCR and Cleanup Kit (5 mg) (T1030 NEB, MA) and validated on 1% agarose gel electrophoresis (Fig. S3). All synthetic DNA gBlocks and PCR fragments were quantified on a Qubit fluorimeter (Invitrogen, OR). The correct sequences of PCR products were validated by Sanger sequencing using an ABI 373 instrument. All restriction endonuclease enzymes, DNA substrates, DNA and protein markers were from New England Biolabs (Ipswich, MA). Sequences of PCR primers and synthetic DNA “gBlocks” encoding putative RE genes are listed in supplementary Table S7.

## Data availability

All data related to the analyses in this work are available from github (https://github.com/nebiolabs/domainator_examples) and Zenodo (https://doi.org/10.5281/zenodo.10989173)

## Code availability

The source code for Domainator is available from https://github.com/nebiolabs/domainator. Scripts for reproducing the analyses presented in this work are available from https://github.com/nebiolabs/domainator_examples

## Supplemental materials

**Supplemental_figures.pdf**: Supplemental figures S1-S6

**Supplemental_tables.xlsx**: Supplemental tables S1-S7

## Acknowledgements

Yu-Cheng Lin for writing a script that became the precursor to Domainator. The many users and beta testers from the research department at New England Biolabs. Gary Smith and the NEB IT team for help with computing infrastructure. The authors are grateful for support from New England Biolabs (NEB), without which this work would not have been possible. The authors are employees of NEB, a manufacturer and vendor of molecular biology reagents. This affiliation does not affect the authors’ impartiality, adherence to journal standards and policies, or availability of data and software.

## Supplementary Figures

**Supplemental Fig S1.**
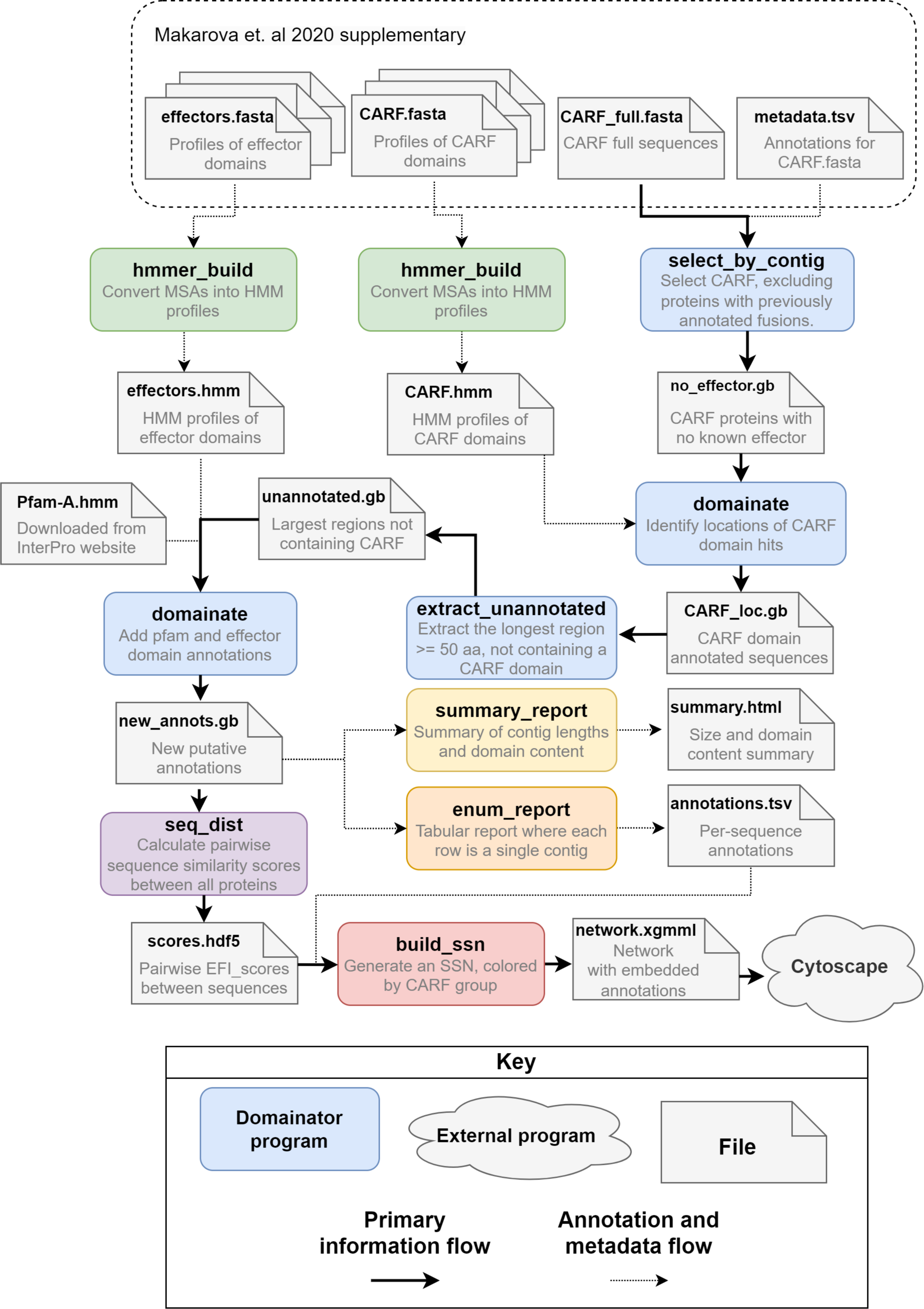
Domainator workflow for analyzing CARF effector domains.

**Supplemental Fig S2.**
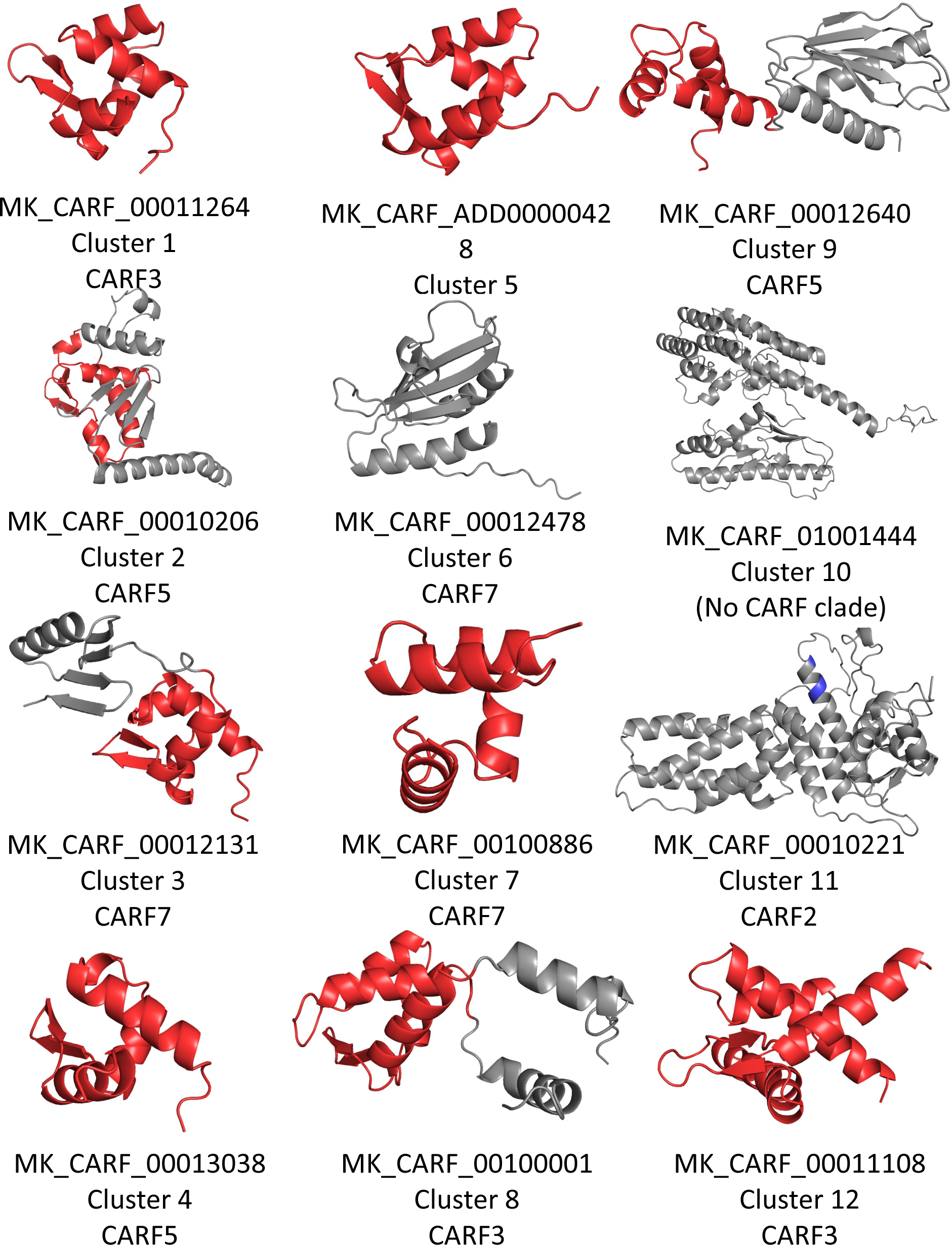
AlphaFold2 predicted structures of unannotated (non-CARF) domains of randomly selected members of each of the 12 largest clusters. Apparent Helix-Turn-Helix or Winged Helix-Turn-Helix DNA binding motifs highlighted in red. Conserved residues of HEPN RNAse RNXXXH highlighted in blue.

**Supplemental Fig S3.**
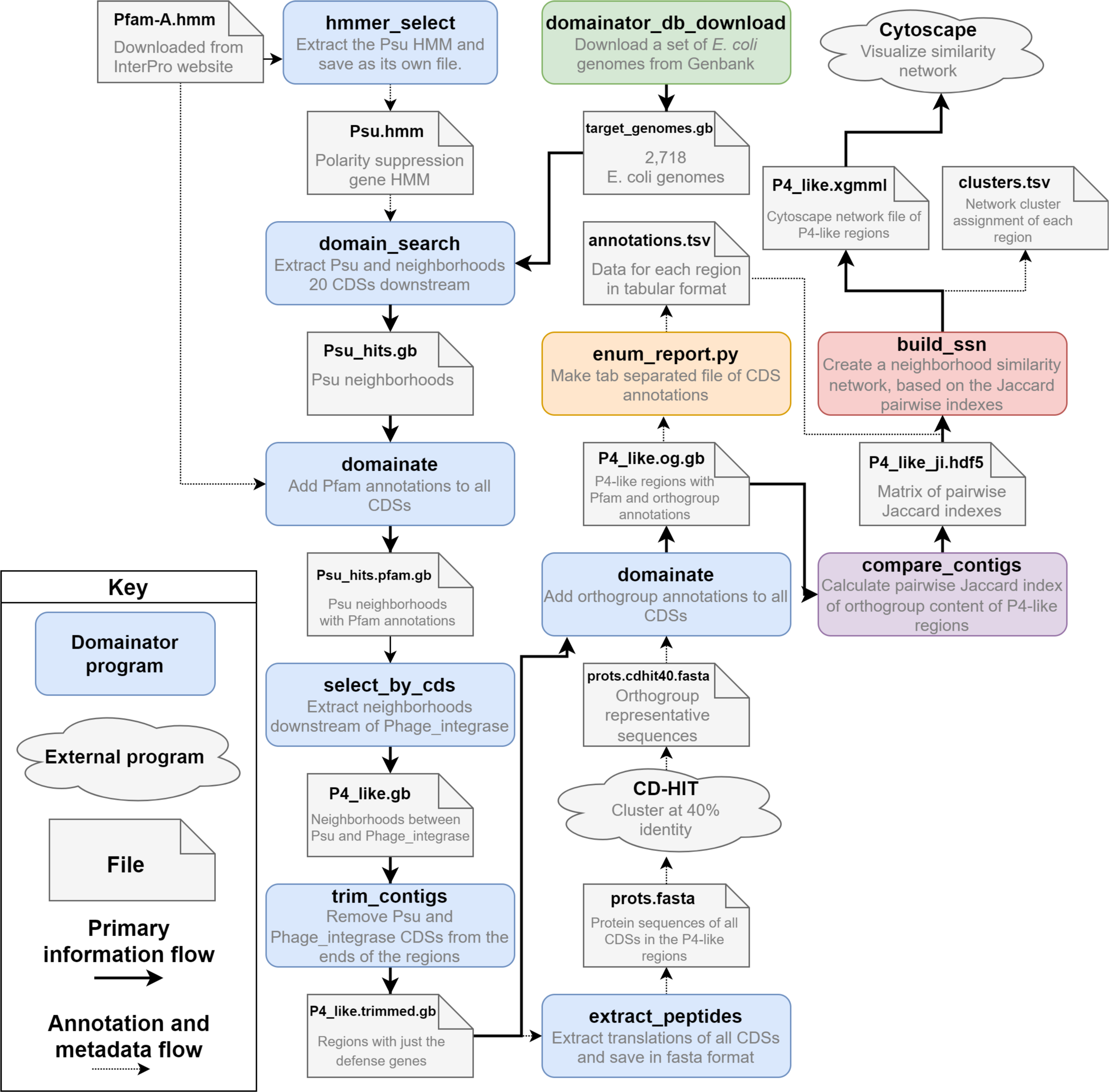
Domainator workflow for analyzing P4-like phage satellite defense hotspots in *E. coli*.

**Supplemental Fig S4.**
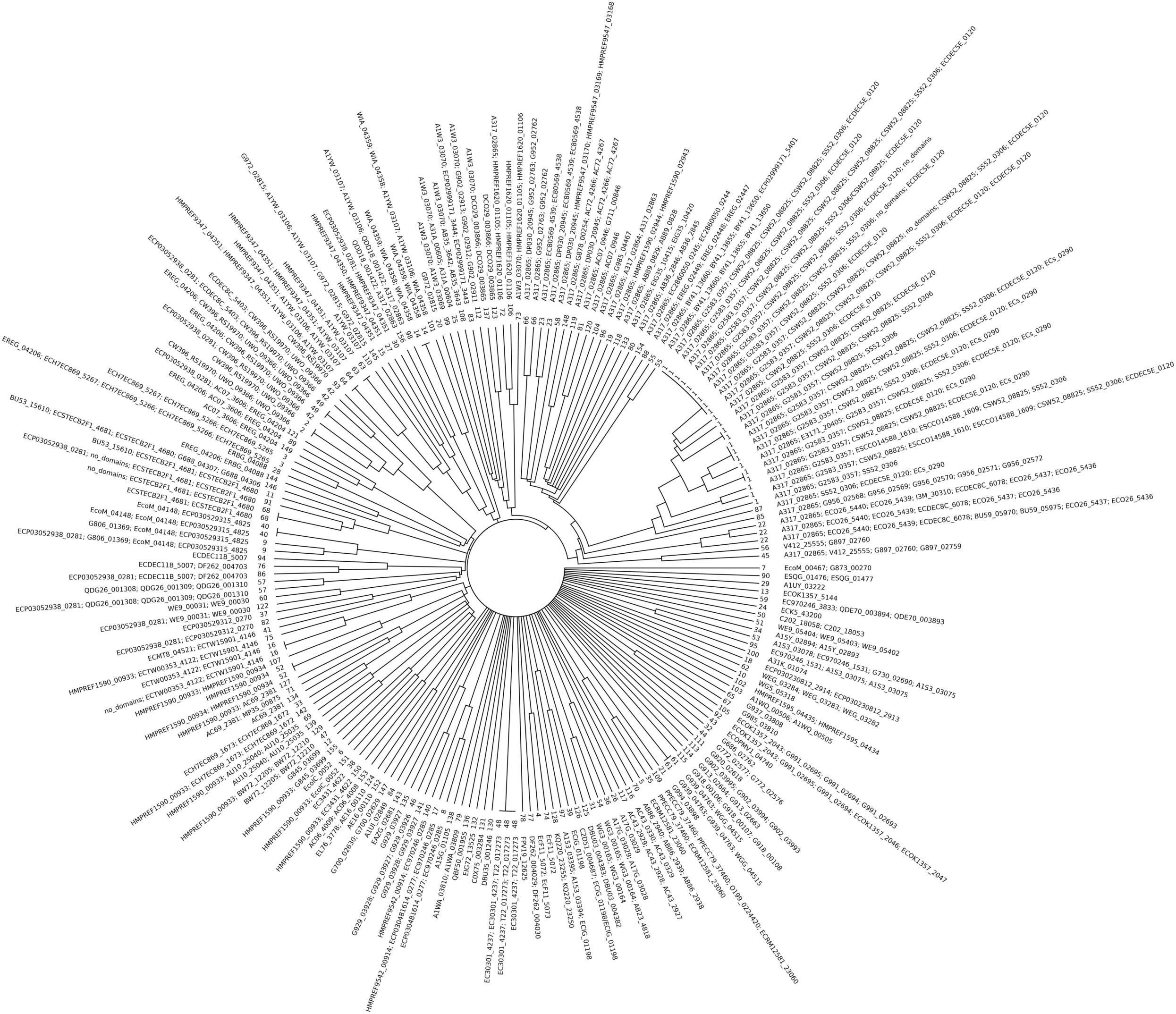
UPGMA tree of Jaccard indexes of orthogroup contents of P4-like satellite defense hotspots. Each leaf is a unique orthogroup architecture. The inner number is the cluster number assignment of the architecture, the outer text is the orthogroup architecture, orthogroup contents of individual CDSs are separated by semicolons “;”, starting with the CDS closest to Psu.

**Supplemental Fig S5.**
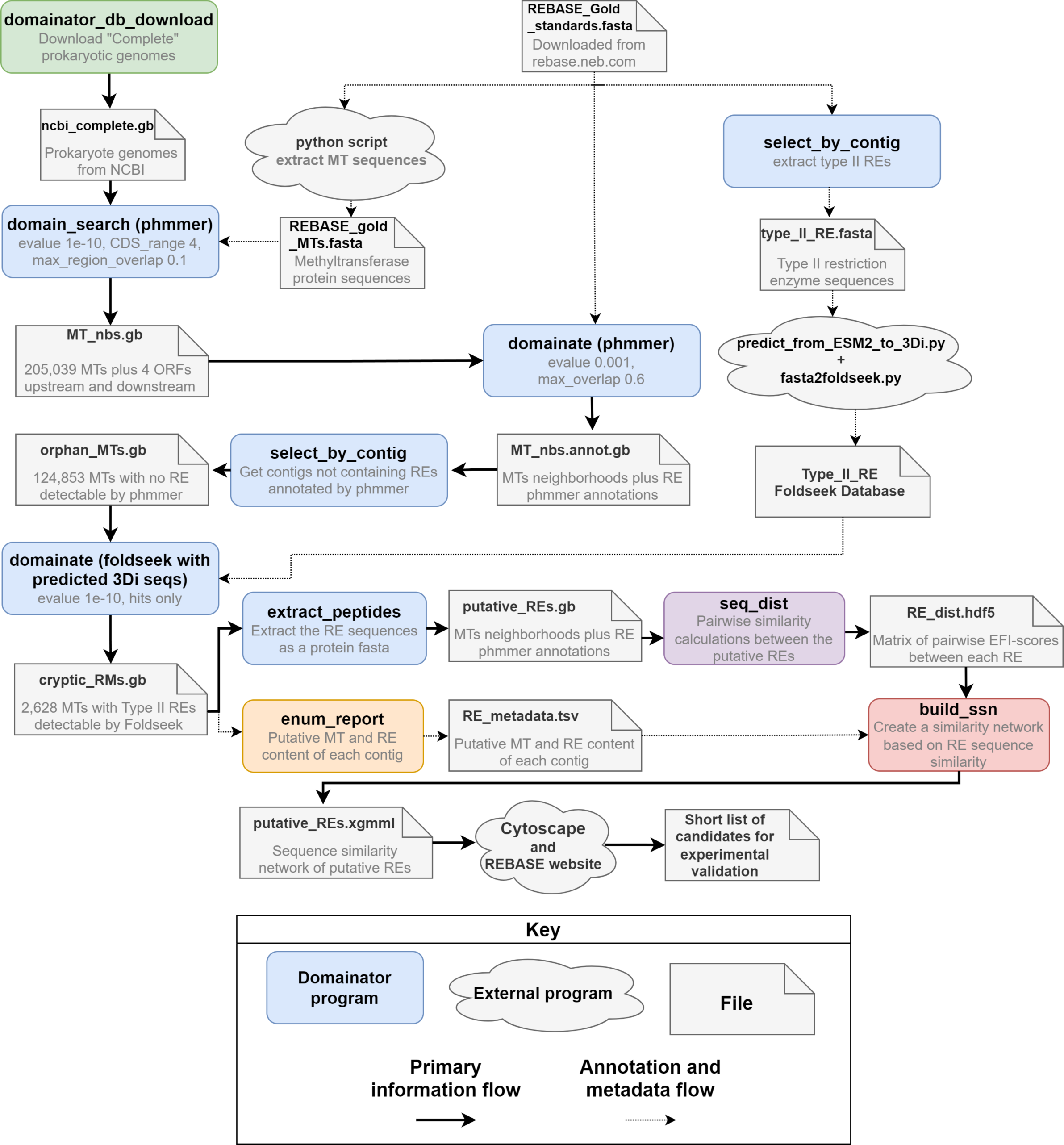
Domainator workflow for identifying cryptic restriction enzymes.

**Supplemental Fig S6.**
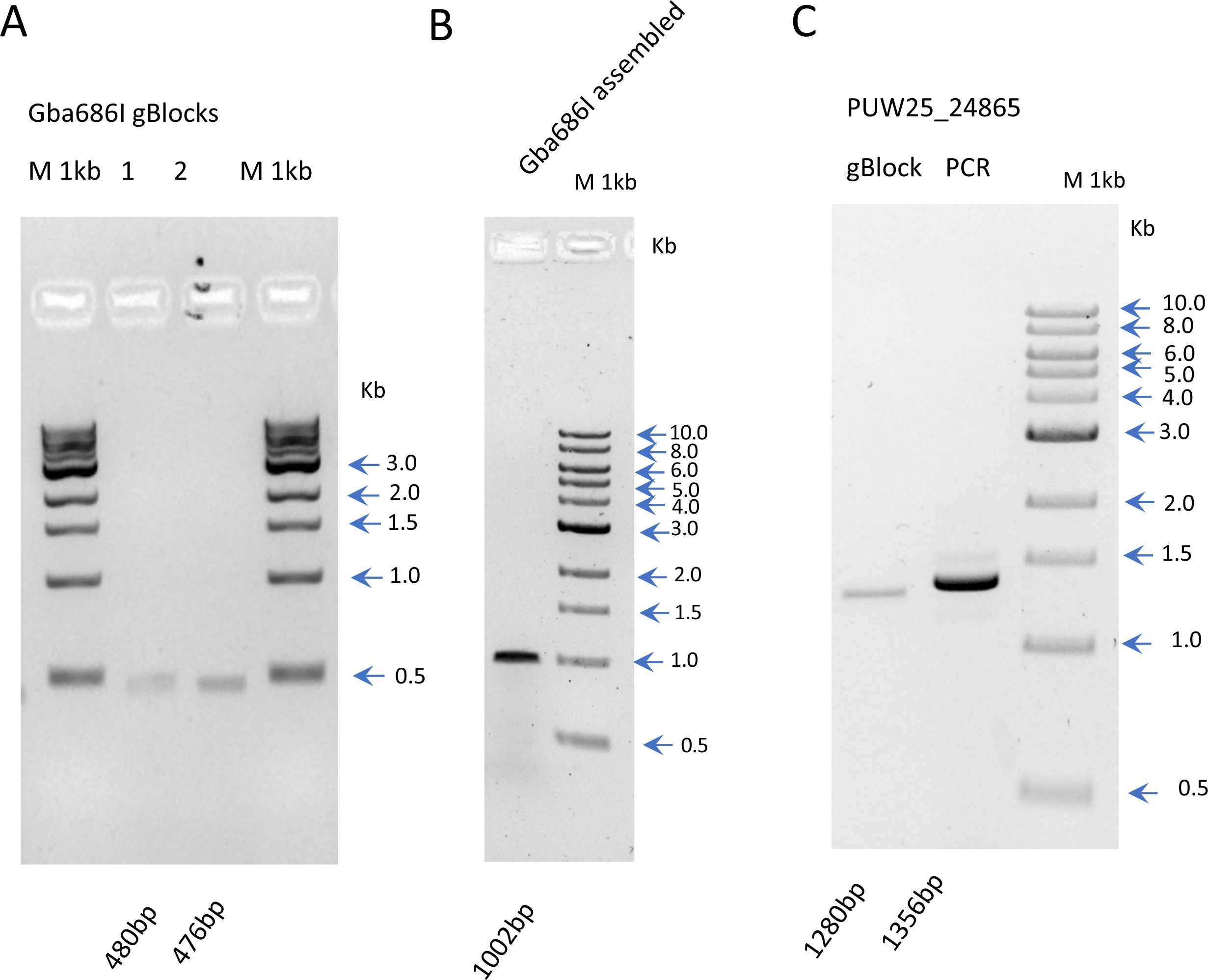
Quality and a size analysis of synthetic gBlocks from IDT and T7 PCR templates for in vitro expression on 1% agarose gel electrophoresis. (A) two gBlocks for Gba686I gene with expected size of 480 and 476 bp. (B) PCR amplified T7 template of Gba686I after assembly with expected size of 1002bp. (C) One gBlock of PUW25_24865 with expected size of 1280bp and PCR amplified T7 template after assembly with expected size of 1356bp.

## References

1. Jumper, J. et al. Highly accurate protein structure prediction with AlphaFold. Nature 596, 583– 589 (2021).

2. Lin, Z. et al. Language models of protein sequences at the scale of evolution enable accurate structure prediction. 2022.07.20.500902 Preprint at 10.1101/2022.07.20.500902 (2022).

3. De Crécy-lagard, V. et al. A roadmap for the functional annotation of protein families: a community perspective. Database 2022, baac062 (2022).

4. Altschul, S. F. et al. Gapped BLAST and PSI-BLAST: a new generation of protein database search programs. Nucleic Acids Res. 25, 3389–3402 (1997).

5. Devos, D. & Valencia, A. Intrinsic errors in genome annotation. Trends Genet. 17, 429–431 (2001).

6. Eddy, S. R. Accelerated Profile HMM Searches. PLOS Comput. Biol. 7, e1002195 (2011).

7. Mistry, J. et al. Pfam: The protein families database in 2021. Nucleic Acids Res. 49, D412–D419 (2021).

8. Aramaki, T. et al. KofamKOALA: KEGG Ortholog assignment based on profile HMM and adaptive score threshold. Bioinformatics 36, 2251–2252 (2020).

9. Cantarel, B. L. et al. The Carbohydrate-Active EnZymes database (CAZy): an expert resource for Glycogenomics. Nucleic Acids Res. 37, D233–D238 (2009).

10. Tatusova, T. et al. NCBI prokaryotic genome annotation pipeline. Nucleic Acids Res. 44, 6614–6624 (2016).

11. Ruhe, Z. C., Low, D. A. & Hayes, C. S. Polymorphic Toxins and Their Immunity Proteins: Diversity, Evolution, and Mechanisms of Delivery. Annu. Rev. Microbiol. 74, 497–520 (2020).

12. Lutz, T. et al. A protein architecture guided screen for modification dependent restriction endonucleases. Nucleic Acids Res. 47, 9761–9776 (2019).

13. Oberg, N., Zallot, R. & Gerlt, J. A. EFI-EST, EFI-GNT, and EFI-CGFP: Enzyme Function Initiative (EFI) Web Resource for Genomic Enzymology Tools. J. Mol. Biol. 435, 168018 (2023).

14. Blin, K. et al. antiSMASH 6.0: improving cluster detection and comparison capabilities. Nucleic Acids Res. 49, W29–W35 (2021).

15. Sibley, M. H. & Raleigh, E. A. Cassette-like variation of restriction enzyme genes in Escherichia coli C and relatives. Nucleic Acids Res. 32, 522–534 (2004).

16. Rousset, F. et al. Phages and their satellites encode hotspots of antiviral systems. Cell Host Microbe 30, 740–753.e5 (2022).

17. Holm, L. Dali server: structural unification of protein families. Nucleic Acids Res. 50, W210–W215 (2022).

18. van Kempen, M. et al. Fast and accurate protein structure search with Foldseek. Nat. Biotechnol. (2023) doi:10.1038/s41587-023-01773-0.

19. Varadi, M. et al. AlphaFold Protein Structure Database: massively expanding the structural coverage of protein-sequence space with high-accuracy models. Nucleic Acids Res. 50, D439–D444 (2022).

20. Ayoub, R. & Lee, Y. RUPEE: A fast and accurate purely geometric protein structure search. PLOS ONE 14, e0213712 (2019).

21. Heinzinger, M., et al. ProstT5: Bilingual Language Model for Protein Sequence and Structure. bioRxiv 2023.07.23.550085 (2023) doi:10.1101/2023.07.23.550085.

22. Johnson, S. R., Peshwa, M. & Sun, Z. Sensitive remote homology search by local alignment of small positional embeddings from protein language models. eLife 12, RP91415 (2024).

23. Makarova, K. S., Anantharaman, V., Grishin, N. V., Koonin, E. V. & Aravind, L. CARF and WYL domains: ligand-binding regulators of prokaryotic defense systems. Front. Genet. 5, (2014).

24. Makarova, K. S. et al. Evolutionary and functional classification of the CARF domain superfamily, key sensors in prokaryotic antivirus defense. Nucleic Acids Res. 48, 8828–8847 (2020).

25. Zallot, R., Oberg, N. & Gerlt, J. A. The EFI Web Resource for Genomic Enzymology Tools: Leveraging Protein, Genome, and Metagenome Databases to Discover Novel Enzymes and Metabolic Pathways. Biochemistry 58, 4169–4182 (2019).

26. Navarro-Muñoz, J. C. et al. A computational framework to explore large-scale biosynthetic diversity from large-scale genomic data. Nat. Chem. Biol. 16, 60–68 (2020).

27. Néron, B. et al. MacSyFinder v2: Improved modelling and search engine to identify molecular systems in genomes. Peer Community J. 3, (2023).

28. Sayers, E. W. et al. GenBank. Nucleic Acids Res. 48, D84–D86 (2020).

29. Paysan-Lafosse, T. et al. InterPro in 2022. Nucleic Acids Res. 51, D418–D427 (2023).

30. Roberts, R. J., Vincze, T., Posfai, J. & Macelis, D. REBASE: a database for DNA restriction and modification: enzymes, genes and genomes. Nucleic Acids Res. 51, D629–D630 (2023).

31. Fu, L., Niu, B., Zhu, Z., Wu, S. & Li, W. CD-HIT: accelerated for clustering the next-generation sequencing data. Bioinformatics 28, 3150–3152 (2012).

32. Edgar, R. C. Search and clustering orders of magnitude faster than BLAST. Bioinformatics 26, 2460–2461 (2010).

33. Lin, K., Zhu, L. & Zhang, D.-Y. An initial strategy for comparing proteins at the domain architecture level. Bioinformatics 22, 2081–2086 (2006).

34. Buchfink, B., Reuter, K. & Drost, H.-G. Sensitive protein alignments at tree-of-life scale using DIAMOND. Nat. Methods 18, 366–368 (2021).

35. Söding, J. Protein homology detection by HMM–HMM comparison. Bioinformatics 21, 951–960 (2005).

36. Steinegger, M. et al. HH-suite3 for fast remote homology detection and deep protein annotation. BMC Bioinformatics 20, 473 (2019).

37. Shannon, P. et al. Cytoscape: A Software Environment for Integrated Models of Biomolecular Interaction Networks. Genome Res. 13, 2498–2504 (2003).

38. Zhang, Y. & Skolnick, J. TM-align: a protein structure alignment algorithm based on the TM-score. Nucleic Acids Res. 33, 2302–2309 (2005).

39. Elnaggar, A. et al. ProtTrans: Towards Cracking the Language of Lifes Code Through Self-Supervised Deep Learning and High Performance Computing. IEEE Trans. Pattern Anal. Mach. Intell. 44, 7112–7127 (2021).

40. Bileschi, M. L. et al. Using deep learning to annotate the protein universe. Nat. Biotechnol. 40, 932–937 (2022).

41. Makarova, K. S. et al. Evolutionary classification of CRISPR–Cas systems: a burst of class 2 and derived variants. Nat. Rev. Microbiol. 18, 67–83 (2020).

42. Steens, J. A., Salazar, C. R. P. & Staals, R. H. J. The diverse arsenal of type III CRISPR–Cas-associated CARF and SAVED effectors. Biochem. Soc. Trans. 50, 1353–1364 (2022).

43. Stella, G. & Marraffini, L. Type III CRISPR-Cas: beyond the Cas10 effector complex. Trends Biochem. Sci. 49, 28–37 (2024).

44. Mirdita, M. et al. ColabFold: making protein folding accessible to all. Nat. Methods 19, 679–682 (2022).

45. Pillon, M. C., Gordon, J., Frazier, M. N. & Stanley, R. E. HEPN RNases – An Emerging Class of Functionally Distinct RNA Processing and Degradation Enzymes. Crit. Rev. Biochem. Mol. Biol. 56, 88–108 (2021).

46. Kita, K. et al. Evidence of Horizontal Transfer of theEcoO109I Restriction-Modification Gene to Escherichia coli Chromosomal DNA. J. Bacteriol. 181, 6822–6827 (1999).

47. Doron, S. et al. Systematic discovery of antiphage defense systems in the microbial pangenome. Science 359, eaar4120 (2018).

48. Cheng, R. et al. A nucleotide-sensing endonuclease from the Gabija bacterial defense system. Nucleic Acids Res. 49, 5216–5229 (2021).

49. Millman, A. et al. An expanded arsenal of immune systems that protect bacteria from phages. Cell Host Microbe 30, 1556–1569.e5 (2022).

50. Jaskólska, M., Adams, D. W. & Blokesch, M. Two defence systems eliminate plasmids from seventh pandemic Vibrio cholerae. Nature 604, 323–329 (2022).

51. Payne, L. J. et al. Identification and classification of antiviral defence systems in bacteria and archaea with PADLOC reveals new system types. Nucleic Acids Res. 49, 10868–10878 (2021).

52. Loenen, W. A. M., Dryden, D. T. F., Raleigh, E. A., Wilson, G. G. & Murray, N. E. Highlights of the DNA cutters: a short history of the restriction enzymes. Nucleic Acids Res. 42, 3–19 (2014).

53. Card, C. O. et al. Cloning and characterization of the HpaII methylase gene. Nucleic Acids Res. 18, 1377–1383 (1990).

54. Camacho, C. et al. BLAST+: architecture and applications. BMC Bioinformatics 10, 421 (2009).

55. Roberts, R. J. et al. A nomenclature for restriction enzymes, DNA methyltransferases, homing endonucleases and their genes. Nucleic Acids Res. 31, 1805–1812 (2003).

56. Zheng, J. et al. dbCAN3: automated carbohydrate-active enzyme and substrate annotation. Nucleic Acids Res. 51, W115–W121 (2023).

57. Tesson, F. et al. Systematic and quantitative view of the antiviral arsenal of prokaryotes. Nat. Commun. 13, 2561 (2022).

58. Cock, P. J. A. et al. Biopython: freely available Python tools for computational molecular biology and bioinformatics. Bioinformatics 25, 1422–1423 (2009).

59. Larralde, M. & Zeller, G. PyHMMER: A Python library binding to HMMER for efficient sequence analysis. Bioinformatics btad214 (2023) doi:10.1093/bioinformatics/btad214.

60. Harris, C. R. et al. Array programming with NumPy. Nature 585, 357–362 (2020).

61. Virtanen, P. et al. SciPy 1.0: fundamental algorithms for scientific computing in Python. Nat. Methods 17, 261–272 (2020).

62. McKinney, W. Data Structures for Statistical Computing in Python. in 56–61 (Austin, Texas, 2010). doi:10.25080/Majora-92bf1922-00a.

63. Hyatt, D. et al. Prodigal: prokaryotic gene recognition and translation initiation site identification. BMC Bioinformatics 11, 119 (2010).

64. Larralde, M. Pyrodigal: Python bindings and interface to Prodigal, an efficient method for gene prediction in prokaryotes. J. Open Source Softw. 7, 4296 (2022).

65. Gerlt, J. A. et al. Enzyme Function Initiative-Enzyme Similarity Tool (EFI-EST): A web tool for generating protein sequence similarity networks. Biochim. Biophys. Acta BBA - Proteins Proteomics 1854, 1019–1037 (2015).

66. Schrödinger, LLC. The PyMOL Molecular Graphics System, Version 2.5. (2015).

